# Lipid Mediated Formation of Antiparallel Aggregates in Cerebral Amyloid Angiopathy

**DOI:** 10.1101/2025.04.21.649887

**Authors:** Ana Pacheco de Oliveira, Divya Baghel, Brooke Holcombe, William Chase, Tyler Ward, Shih-Hsiu J. Wang, Ayanjeet Ghosh

**Affiliations:** Department of Chemistry and Biochemistry, The University of Alabama, Tuscaloosa, AL 35401, USA; Departments of Pathology and Neurology, Duke University, Durham, NC 27710, USA

## Abstract

Cerebral amyloid angiopathy (CAA) is a cerebrovascular disorder marked by amyloid-β (Aβ) deposition in blood vessel walls, leading to hemorrhage and recurring stroke. Despite significant overlap with Alzheimer’s disease (AD) through shared Aβ pathology, the specific structural characteristics of Aβ aggregates in CAA and their variations between stages of disease severity are yet to be fully understood. Traditional approaches relying on brain-derived fibrils can potentially overlook the polymorphic heterogeneity and chemical associations within vascular amyloids. This study utilizes sub-diffraction, label-free mid-infrared photothermal (MIP) spectroscopic imaging to directly probe the chemical structure and heterogeneity of vascular amyloid aggregates within human brain tissues across different CAA stages. Our results demonstrate a clear increase in β-sheet content within vascular Aβ deposits corresponding to disease progression. Crucially, we identify a significant presence of antiparallel β-sheet structures, particularly prevalent in moderate/severe CAA. The abundance of antiparallel structures correlates strongly with co-localized lipids, implicating a lipid-mediated aggregation mechanism. We substantiate the *ex-vivo* observations using nanoscale AFM-IR spectroscopy and demonstrate that Aβ40 aggregated *in vitro* with brain-derived lipids adopts antiparallel structural distributions mirroring those found in CAA vascular lesions. This work provides critical insights into the structural distributions of Aβ aggregates in CAA, highlighting the presence of polymorphs typically associated with transient intermediates, which may lead to alternate mechanisms for neurotoxicity.

## Introduction

Cerebral amyloid angiopathy (CAA) is a cerebrovascular disorder that involves accumulation of the amyloid beta (Aβ) peptide in cerebral blood vessels, which leads to compromised vascular integrity resulting in hemorrhage and recurring strokes(1–3). Currently there are no treatments available for CAA, and definitive diagnosis of CAA requires postmortem examination of the brain. There is significant overlap between CAA and Alzheimer’s disease (AD), and an estimated 70% of AD patients exhibit hallmarks of CAA pathology(4, 5). On a molecular level, this is expected since both CAA and AD involve aggregation of the amyloid beta (Aβ) peptide, which is the primary component of both amyloid plaques found in AD and the vascular deposits characteristic of CAA(3, 5, 6). While the deposition of Aβ is common to both AD and CAA, their pathological pathways diverge in terms of clinical manifestation and mechanism of neurodegeneration. The aggregation of Aβ has been studied in depth, both *in-vitro* and *ex-vivo*; and it is well established that Aβ polymorphs, whether formed *in-vitro* or isolated from the brain, largely adopt a parallel cross β structure(7–16). However, it is not fully understood if these structural insights are equally applicable to AD and CAA. Thus, while significant progress has been made in understanding CAA, fundamental questions still persist regarding its pathogenesis and progression. In comparison to AD, less is known about the fibrillar structures relevant to CAA, and recent cryo-Electron Microscopy (cryo-EM) studies have shown that the structures of Aβ fibrils isolated from plaques are distinct from that of Aβ40 filaments in CAA, suggesting that AD and CAA involve different pathogenic species, albeit arising from the same peptide(17–21). A key challenge in understanding CAA specific Aβ polymorphs is probing the structure and chemistry of the Aβ aggregates in the brain. While *in-vitro* structural models have revealed fundamental insights into amyloid structure, it is not known how translatable these structures are to *in-vivo* amyloid deposits. The structure of amyloid aggregates in the brain, for both parenchymal plaques and vascular deposits, are not well understood. While recent cryo-EM studies of brain derived fibrils have addressed this to some extent, the variations that might persist between different pathological lesions in terms of amyloid structure are not always apparent in such approaches. As a result, the relationship between the specific aggregate structures of Aβ and their pathogenicity remains unclear. Furthermore, it is not known if there are variations in the structure and chemical associations of these aggregates across different disease stages. Additionally, Aβ aggregates in brain lesions can exhibit considerable heterogeneity in terms of composition. While Aβ aggregates are the major component of plaques, concurrent presence of other proteins such as tau, synuclein and apolipoprotein E (apoE) colocalized with Aβ has been conclusively demonstrated(22). Furthermore, multiple isoforms of Aβ can be present in brain amyloid aggregates(23). Recent studies have evidenced potential heterotypic interactions between different Aβ isoforms and other brain proteins leading to altered fibrillar structures(24). ApoE is known to associate with Aβ and alter the structure of early stage Aβ aggregates, leading to their impaired clearance from tissues.(25–27) Tau, on the other hand, is known to suppress the fibrillization of Aβ, essentially locking it in a oligomeric state(28). Lipid mediated aggregation pathways can also be particularly relevant for vascular aggregates, where interaction between Aβ and the cellular components in the tunica media can modulate the kinetics and structure of aggregates. However, the applicability of these to CAA is yet to be determined and adds to the complexity of its molecular pathology. The composition of vascular aggregates and heterogeneities therein at different disease stages may have significant implications in the context of pathological manifestations and disease progression.

The critical gap in knowledge of amyloid structures in CAA brain stems largely from limitations of structural biology techniques such as Solid State Nuclear Magnetic Resonance (ssNMR) and cryo-EM towards direct tissue measurements. Structural assessments using these approaches typically rely on characterizing fibrils amplified from seeds derived from diseased brains. However, it is not fully known if brain derived fibrils adequately represent the polymorphic diversity and structural heterogeneity that may exist between different amyloid deposits. Moreover, information on association of amyloid aggregates with other chemical moieties of interest are lost when characterizing brain derived fibrils with *in-vitro* techniques. Recent ssNMR studies have demonstrated the possibility of antiparallel structures in CAA derived fibrils, which are not observed in later generations in seeded growth(29). This is in contrast to structures derived from cryo-EM, and these seemingly contradictory findings highlight the likely heterogeneity underlying vascular aggregates and underscore the need for structural measurements of the same directly in tissues(18). Transient antiparallel structures have been shown to exist for Aβ mutants(30, 31), and also in in amyloid plaques in AD(32), but their role in CAA is yet to be understood. Determination of the prevalence of such intermediates in vascular amyloid deposits and structural assessment of vascular aggregates in heterogeneous tissue environments necessitate integration of spectroscopies, which offer structural sensitivity, with spatial resolution, so that facets of individual deposits can be probed.

Infrared (IR) spectroscopic imaging, which augments the structural insights of IR with microscopy(33–35) is an experimental approach uniquely suited for this challenge and has been applied towards studying amyloid plaques and heterogeneities thereof(32, 36–42). However, applications of IR imaging towards characterization of the chemistry of vascular aggregates in tissues has remained unrealized so far. In this work, we employ mid infrared photothermal (MIP) spectroscopic imaging to investigate the chemical heterogeneities of vascular amyloid aggregates found in diseased human brain tissues from different CAA stages. MIP microscopy is a recently developed technique that improves the spatial resolution of conventional IR imaging by approximately an order of magnitude by leveraging thermal lensing arising from photothermal response of the specimen to indirectly measure the IR absorption(43–49). MIP thus provides the optimal combination of high resolution and label-free imaging capabilities, making it ideal for mapping the chemistry of vascular amyloids. Using MIP imaging, we demonstrate that vascular aggregates in CAA exhibit increase in β-sheet content with disease stages. We further show significant presence of antiparallel structures in vascular amyloids, particularly in moderate/severe CAA, which is correlated to the abundance of colocalized lipids and thus indicative of a lipid-mediated aggregation mechanism. Finally, to get molecular insights into the origin of these observations, we use AFM-IR, which couples IR spectroscopy to Atomic Force Microscopy (AFM) and show that the structural distribution of Aβ40 fibrils aggregated in presence of brain derived lipids mirrors the trends observed in vascular aggregate, thus proving a molecular basis for disease relevant pathogenic polymorphs.

## Materials and Methods

### AD Tissue Samples

Formalin fixed paraffin embedded (FFPE) tissue sections of the human mid frontal cortex were obtained from the Bryan Brain Bank of the Duke-UNC Alzheimer’s Disease Research Center (ADRC). Postmortem evaluation of AD neuropathologic change and CAA severity was performed by a pathologist at the Bryan Brain Bank following established guidelines(50, 51) prior to any analysis. The details of the specimens are provided in the Supporting Information (Table S1). For each specimen, two parallel sections were used. One tissue section was immunohistochemically (IHC) stained with beta-amyloid antibody 4G8 (BioLegend Cat#: 800702; 1:1000) for identification of the vascular amyloid deposits and other amyloid features at the Bryan Brain Bank. The other section was used for MIP microscopic analysis. The section used in the MIP microspectroscopy analysis was mounted on an IR reflective slide (MirrIR Low-E, Kevley Technologies) to minimize any spectral artifacts from the substrate. Both tissue samples were sectioned to ∼5 µm in thickness. Brightfield images of the IHC stained tissue were acquired using an Olympus BX43 at 40x magnification using manual WSI software (Microvisioneer) for imaging stitching. The brightfield images of the IHC stained tissue sections were used as a reference to locate the vascular amyloid deposits. The tissue sections on the Low-E slide were deparaffinized by submerging the tissue sections in n-hexane for 24 hours followed by storage under mild vacuum until the MIP analysis.

### Photothermal Infrared Microscopy

A photothermal microscope (mIRage, Photothermal Spectroscopy Corp.) equipped with a tunable quantum cascade infrared (IR) laser (MIRcat, Daylight Solutions) and 532 nm detection laser was used for the MIP measurements. All MIP spectra and discrete frequency images were acquired using a 0.78 NA 40x Cassegrain objective. A range of 7-12 spectra were collected in the inner walls of the vascular amyloid deposits while 9-15 spectra were collected in the immediate area surrounding the vascular amyloid deposits. Discrete frequency images of specific wavenumbers were collected using the same parameters as the spectra, at 0.5 µm resolution and 50 µm /pt scanning rate.

### Data Processing

All data collected in this study was processed using MATLAB. The MIP spectral data was baseline corrected by subtracting a linear background. All spectra were smoothed using a 3-point moving average and (2,7) Savitzky-Golay filter. The second derivative spectra were smoothed using a (3,11) Savitzky-Golay filter. The discrete frequency ratio images were denoised using a two-point median filter. Ratio images were created by a ratio of two peaks of interest within the Amide-I region that correspond to relevant signatures of known protein secondary structures. These ratio images provide a method for normalization and visualization between local areas of analysis and the entire tissue sample to account for natural variations in density and thickness that occur across the overall sample area. The following ratios were used: 1628:1660 and 1692:1660. These ratios represent parallel and antiparallel β-sheet, respectively. Multivariate Curve Resolution with Alternating Least Squares (MCR-ALS) was used to identify spectral components with 4 initial components, 4 non-negative constraints, fast nonnegative least squares(fnnls) implementation, and 500 iterations. AFM images were processed by Gwyddion software. IR data were analyzed by using MATLAB software by applying (3, 7) Savitzky-Golay filter and a 5-point moving average filter. A baseline correction was applied for each spectrum. The statistical analysis of the spectra was performed in Python, using the Pingouin package.

### Aggregation of 13C**IZI**labeled A**β**-40

^13^C-Aβ40 (rPeptide, USA) was first treated with 1,1,1,3,3,3-hexafluoroisopropanol (HFIP) to destroy any preformed aggregates. HFIP was completely evaporated at room temperature (24 °C), under vacuum. 100 μg of ^13^C-Aβ40 was prepared in 10 mM phosphate buffer.

### Coaggregation of 13C**IZI**labeled A**β**40 with Brain Polar Lipid Extract

A 25 mg/mL solution of brain polar lipid extract (Avanti Polar Lipids) in chloroform (CHCl_3_) was used. A 20 μL aliquot was transferred to a separate vial and placed in a vacuum desiccator to allow complete evaporation of CHCl_3_. Subsequently, 10 mM phosphate buffer was added, leading to turbidity within a few minutes, indicating the formation of lipid vesicles. The solution was then vortexed for 1 minute and sonicated for 30 seconds before being added with amyloid-beta to prepare the coaggregation mixture. 25 mg/ml brain polar lipid extract was co-aggregated with ^13^C-Aβ40 in 10 mM phosphate buffer, pH = 7.4, in 5:1 molar ratio. The aggregation was performed for 24 hours at 37 °C without any agitation. For control, brain polar lipid extract dissolved in DI water, with similar concentration as in coaggregation mixture was used.

### AFM**IZI**IR Analysis

AFM-IR experiments were carried out by Bruker NanoIR3 instrument equipped with mid-IR quantum cascade laser (MIRcat, Daylight solutions). Experiments were performed at room temperature and relative humidity inside the instrument was kept low by continuous purging with dry air. Both AFM imaging and IR data collection was done in tapping mode with cantilevers having resonance frequency of 75±15 kHz and spring constant of 1-7 N/m. AFM scan rate were varied from 0.5 Hz to 1.0 Hz. First a high-resolution AFM image of the sample was recorded having multiple oligomers/fibrils in the scan area. Then AFM tip was placed on individual oligomer/fibril at random to obtain the IR spectra. For both samples, 35 spectra from two different spatial locations were acquired from oligomer/fibrils to avoid under sampling. IR spectral resolution was 2 cm^-1^. 128 coadditions at each point and 16 co-averages for each spectrum were applied.

## Results and Discussion

### Antiparallel structure in vascular aggregates

In this study, a total of 80 vascular amyloid deposits from one normal and four diseased mid-frontal brain tissue specimens corresponding to different stages of CAA severity were analyzed. Typically, CAA severity is graded according to the degree of amyloid deposition in the blood vessel walls and resulting pathological features. In mild CAA, blood vessels show focal limited amyloid deposition without vascular damage, whereas moderate and severe CAA exhibit more pronounced circumferential amyloid presence with associated vascular damage, particularly for the latter(3). Amyloid positive blood vessels were identified using immunohistochemical (IHC) staining and grouped into two categories: mild CAA and moderate/severe CAA based on the extent of amyloid deposition. Figure 1 shows a representative example of each IHC stained vascular aggregate type. Additional images are shown in the Supporting Information (Figures S1-S2). From a molecular perspective, it is well known that Aβ fibrils have an ordered parallel β-sheet structural motif, while prefibrillar aggregates can have a comparatively disordered structure. Both parallel and antiparallel β-sheets have unique vibrational signatures in amide I region of mid-infrared spectra, arising from excitonic coupling between the backbone amide moieties. Both exhibit a characteristic band at 1625-1635 cm^-1^, while antiparallel β structures exhibit an additional band at around 1690 cm^-1^(52, 53). Figures 1 E-H show MIP images at 1628 cm^-1^ and 1692 cm^-1^ normalized to the spectral intensity at 1660 cm^-1^, reflective of overall β-sheet and antiparallel β-sheet populations respectively, for the blood vessels in Figure 1 A-B. An unstained, parallel tissue section was used for acquisition of IR data to avoid spectral interferences from the stain. This is a commonly used approach in IR imaging(33, 35) and we have previously used this strategy in our studies targeting Aβ plaques(32, 54, 55). We observe increased MIP intensity from the blood vessel walls for moderate/severe CAA, consistent with elevated presence of β-sheet rich amyloid aggregates. Interestingly, we also observe signatures for antiparallel β-structures from the blood vessel indicating a complex structural ensemble different from typical parallel cross-β fibrils. In comparison, the blood vessel corresponding to mild CAA does not exhibit significantly enhanced parallel or antiparallel β-sheet signatures. Of course, this does not indicate how generalizable this observation is and if such transient structures persist in other vascular aggregates as well. However, expanding the discrete frequency imaging approach above to a statistically relevant number of blood vessels is challenging for two reasons. Firstly, while normalized IR intensities are reflective of relative populations of structural motifs such as β-sheets, they cannot be used to quantitatively assess the overall secondary structure of protein aggregates and changes therein. Furthermore, a discrete frequency approach essentially requires preempting the bands of interest, which is not always viable. Acquisition of the entire amide I spectra and subsequent spectral deconvolution represents a more quantitative alternative, which, however, significantly increases the experimental time. To mitigate this, we therefore employed a spatially resolved spectroscopic approach, wherein full spectra of specifically the blood vessel walls were acquired, as indicated in Figure 2A (green dashed circle). This offers the same spectral insights into the composition of amyloid rich blood vessels. For each blood vessel, 10-30 spectra were acquired, depending on its size. The mean spectra of blood vessels corresponding to mild and moderate/severe CAA thus acquired are shown in Figure 2B. Mean spectra acquired from control blood vessels that are amyloid negative are also shown for comparison. We observe that the mean spectra of vascular amyloids corresponding to moderate/severe CAA exhibit significantly enhanced intensity at ∼ 1632 cm^-1^ compared to normal blood vessels, indicating an increase in β-sheet abundance. The spectra also show a distinct peak at ∼1736 cm^-1^. This absorption is characteristic of carboxylic acids and ester functional groups, and in the context of cellular and tissue imaging, is attributed to lipid moieties(33, 38). The spectra thus provide evidence supporting a concurrent increase in lipids along with β-sheet motifs, which is a unique finding and highlights the possibility of a lipid driven aggregation mechanism underlying the pathological changes in CAA. In addition to the above, we also note an increase in the spectral intensity at ∼ 1686 cm^-1^, which can be assigned to antiparallel β structures. In comparison, vascular aggregates from mild CAA exhibit a relatively smaller increase for both the overall and antiparallel β-sheet populations. An increase in the lipid band is also observed, compared to normal blood vessels. Taken together, the spectra indicate an overall increase in β-sheet population with increasing CAA severity, potentially correlated to lipids, and also raise the possibility of concurrent presence of antiparallel β-structure.

**Figure 1.**
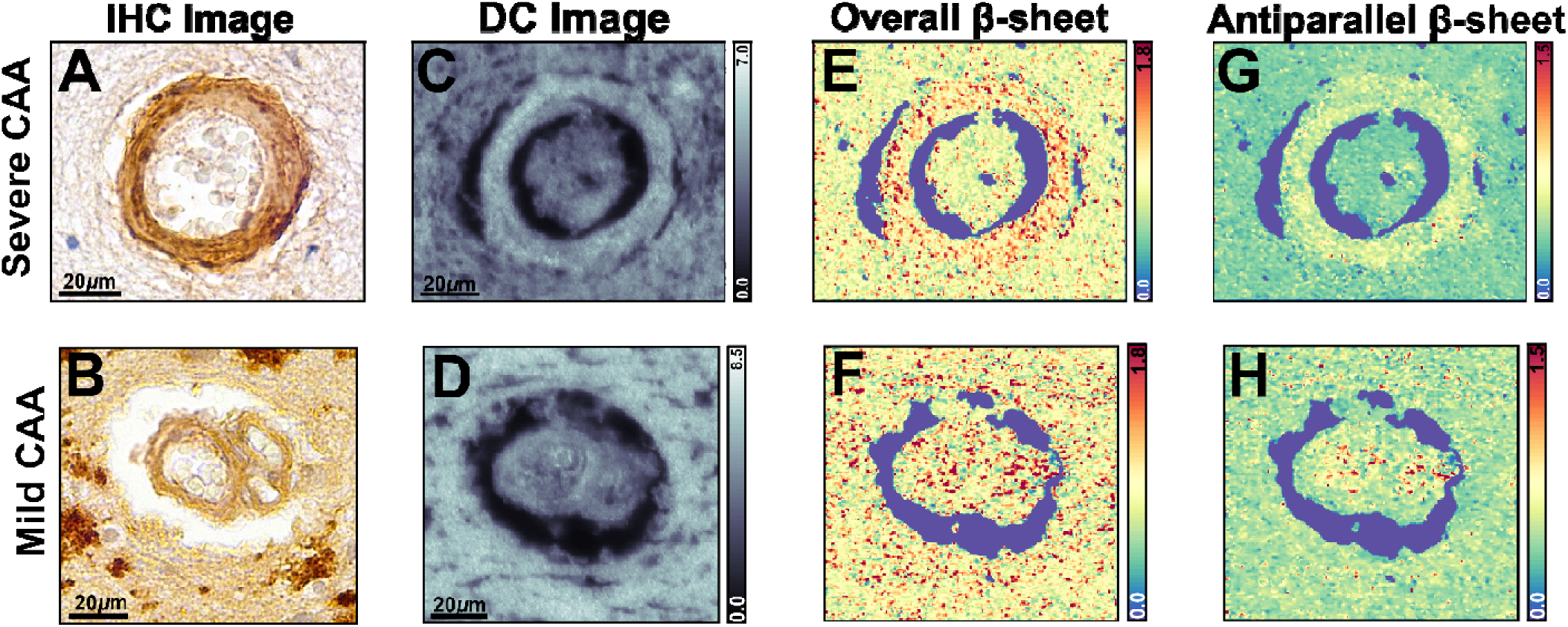
(A, B) Brightfield optical images IHC stained vascular amyloid deposits from a severe and a mild CAA case. (C, D) DC image of vascular amyloid deposits from A and B. (E, F) IR ratio images (_I1628_/I_1660_) highlighting the presence of overall β-sheet content. (G, H) IR ratio images (I_1692_/I_1660_) reflective of antiparallel β-sheet populations in the vascular deposits.

**Figure 2.**
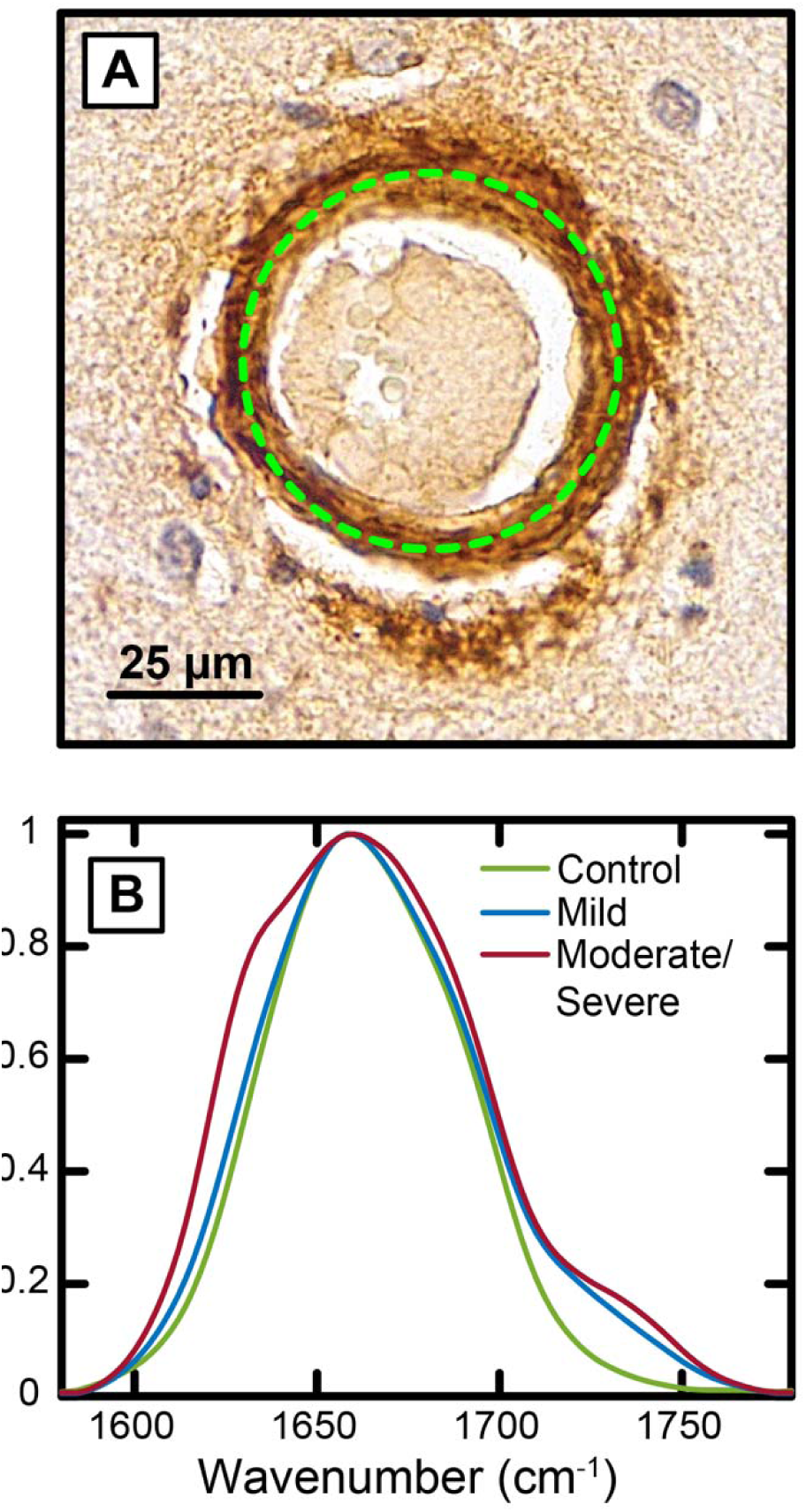
(A) Schematic representation of the spatially resolved spectroscopic approach used for vascular amyloid characterization. The dashed circle represents the locations along the blood vessel wall from which spectra are acquired. (B) Mean IR spectra of vascular aggregates from control, mild and moderate/severe CAA.

### Antiparallel β-structure in CAA is correlated to lipids

The amide I band in proteins is a convolution of multiple underlying components, and as a result, the spectral intensities are not quantitative reflections of the relative populations of different structural motifs. It is thus difficult to unambiguously establish the above conclusions without further analysis. Therefore, to gain further insights into the secondary structure of the vascular amyloids and their evolution with disease severity, we performed spectral deconvolution using Multivariate Curve Resolution-Alternating Least Squares (MCR-ALS). The gold standard in IR spectral deconvolution is band fitting; however, it requires a priori knowledge of the spectral components, such as frequency and linewidth. MCR-ALS is a matrix factorization approach that enables blind deconvolution of spectral data without the need for information of the pure spectra(56–58). By iteratively applying alternating least squares optimization with constraints such as non-negativity and unimodality, MCR-ALS decomposes the data matrix into pure spectral profiles and their corresponding concentration profiles, similar to global fitting of spectra. This allows for quantitative identification of individual constituents and tracking their evolution across datasets, making it an ideal tool for assessing the evolution of secondary structure in CAA, as presented above. The results from MCR-ALS are presented in Figure 3A. The spectra were grouped by blood vessel for this analysis to mitigate any potential effects from size variations. We find that the spectra can be adequately described as a weighted sum of four bands centered at 1628 cm^-1^, 1656 cm^-1^, 1690 cm^-1^ and 1732 cm^-1^, attributable to β-sheets, random coils/disordered structures, antiparallel β-sheets and lipids, respectively. The weights of each component from the different blood vessel groups are shown in Figure 3B. The trends, as determined from MCR-ALS, for each of the spectral components, across the vascular aggregates from different CAA stages are shown in Figure 3 C-F. We observe that the overall β-sheet increases in mild and moderate/severe CAA compared to control. The exact opposite trend is observed for the random coil/disordered population. Both antiparallel β-sheets and lipids also exhibit a trend similar to overall β-structure, increasing steadily from control to mild and moderate/severe CAA. We note the relative changes in secondary structure populations are small, which is somewhat expected in complex tissue environments, where other proteins can also contribute to the overall spectrum. To verify if our observations are statistically significant, we performed Welch’s ANOVA followed by a Games-Howell post-hoc test for each structural component. The p-values, listed in the Supporting Information (Table S2), indicate that there exists statistically significant difference in overall and antiparallel β-sheets and lipids between normal blood vessels and those from moderate/severe CAA. The overall populations for mild CAA are not always significantly different from those observed for the control and moderate/severe case, which likely arises due to a relatively low abundance of Aβ aggregates. We address the implications of potential contribution from non-Aβ protein components to the spectra later in the manuscript. Taken together, the above trends suggest a direct relationship between abundance of β-sheet structures, parallel and antiparallel, and lipids, which in turn implies a correlation between the respective weights or populations of these moieties. The correlation of both overall and antiparallel β-sheet weights with those of lipids, as obtained from MCR-ALS are shown in Figure 4 A-B. We find that lipids correlate with presence of antiparallel β-structures, but not with the overall β-sheet population. This indicates that a lipid mediated aggregation pathway specifically contributes to formation of transient, antiparallel structures. The overall β-sheet abundance is not strongly correlated to lipids likely because of presence of both parallel and antiparallel species, where only the latter is connected to lipids.

**Figure 3.**
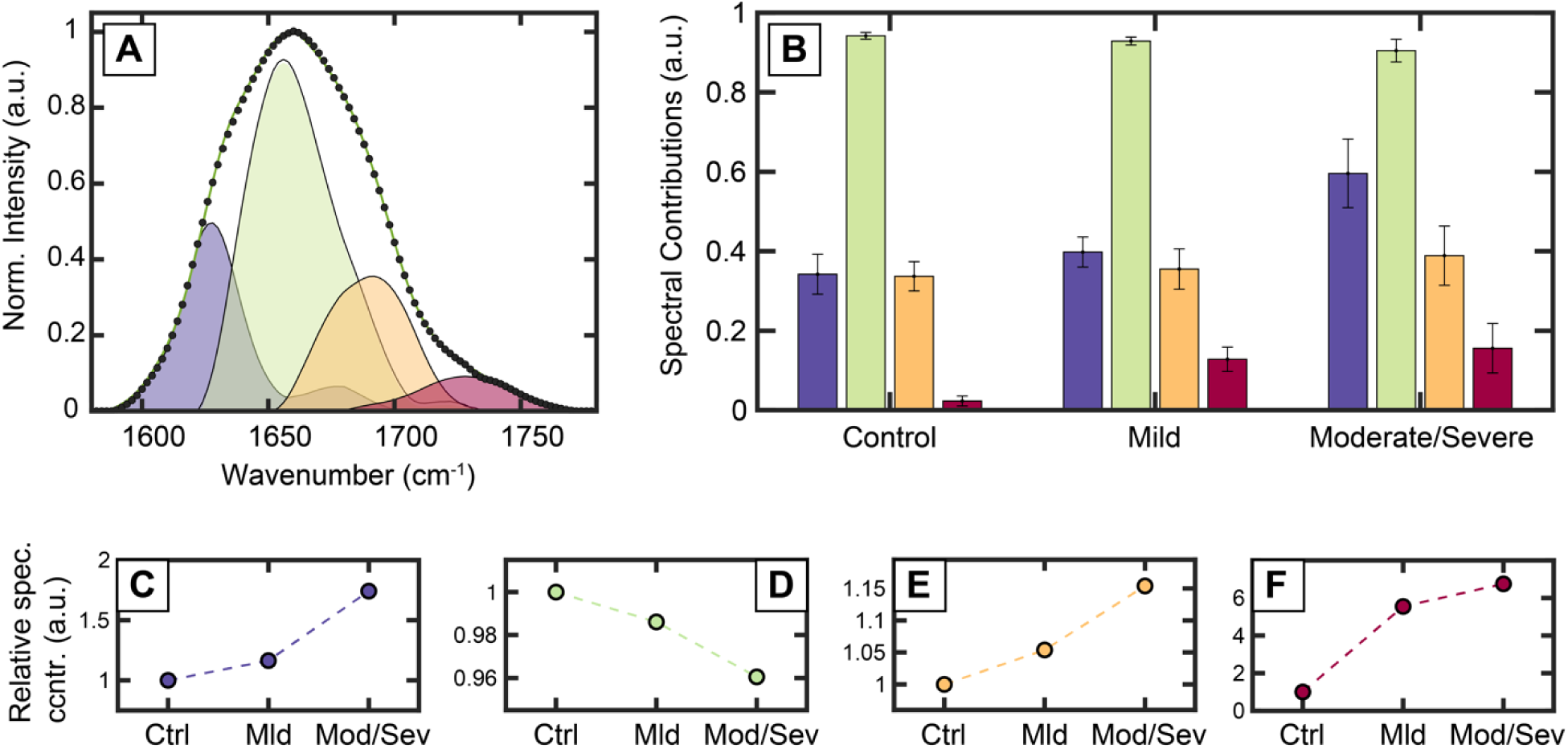
(A) Spectral deconvolution of the amide I IR spectra showing its individual components. Overall β-sheet (purple), antiparallel β-sheet (green), disordered structures (yellow) and lipids (red). (B) Contributions of each spectral component determined from deconvolution across blood vessels from control, mild and moderate/severe CAA. (C-F) The relative change in the populations of each spectral component, shown in B. The values have been normalized to that of the control, for clarity.

**Figure 4.**
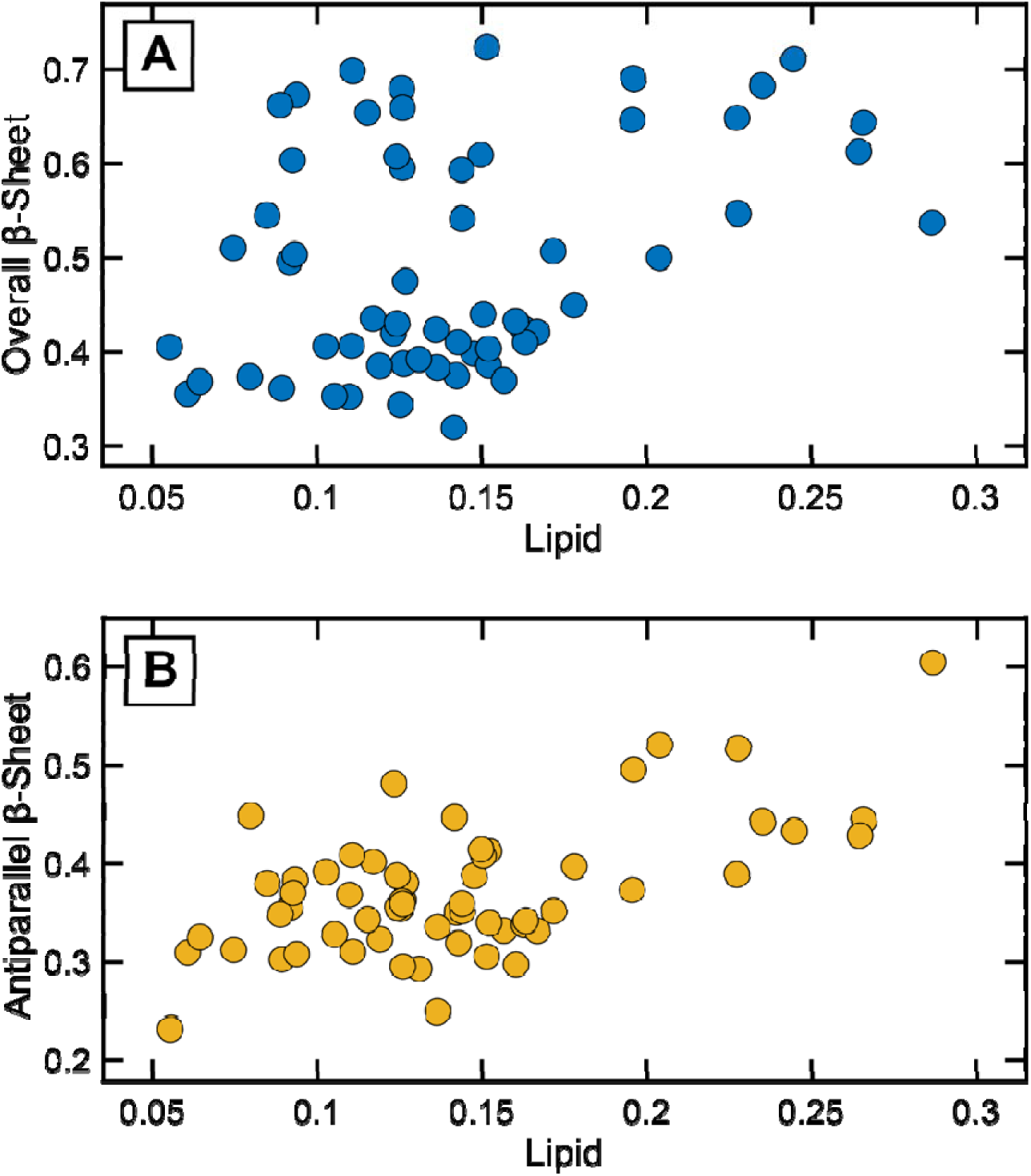
Correlations of lipid populations with β-sheets, as determined from spectral deconvolution. (A) Scatter plot of spectral weights of overall β-sheets versus lipid content. (B) Scatter plot of spectral weights of antiparallel β-sheets versus lipid, exhibiting a more linear correlation, which implies that presence of antiparallel β-sheet structures depends on lipids.

The severity of CAA is linked to increase in fibrillar amyloid aggregates in cerebral vasculature, leading to fragmentation of the blood vessel walls and hemorrhage(3, 5). Our results are consistent with this view: we find gradual increase in the overall β-sheet population, characteristic of amyloid fibrils, going from normal blood vessels to those corresponding to mild and moderate/severe CAA. The presence of antiparallel structures, however, is unexpected. It is well known that Aβ fibrils have a parallel cross β arrangement, whereas oligomers can have an antiparallel structure. The antiparallel structure, however, has also been demonstrated to exist in Aβ mutants, where transient fibrillar intermediates can have this structural motif, while mature fibrils are parallel(30, 59). More recently, the presence of antiparallel structure in wild-type Aβ40, the isoform most relevant to CAA, has also been evidenced(24). The increase in antiparallel population in moderate/severe CAA can thus be attributed to increased presence of either antiparallel fibrils or oligomers, or both. It should be noted that our findings do not point to presence of exclusively antiparallel fibrils in severe/moderate CAA, and it likely that a mix of parallel and antiparallel fibrils exist in these vascular deposits. Furthermore, the presence of only antiparallel structures would lead to an identical evolution for overall β-sheet population between the specimens; we observe that the trends of overall and antiparallel β-sheets, while similar, are not identical. However, presence of antiparallel structures is inconsistent with recent findings from cryo-EM, which have unequivocally demonstrated that the fibrillar polymorphs associated with CAA have a parallel cross-β motif(19, 60). It is important to note in this text that cryo-EM studies of brain derived fibrils usually do not focus on fibrillar aggregates from individual blood vessels or plaques. Furthermore, it is not fully understood how different polymorphs isolated from diseased brains propagate relative to each other in seeded growth. Therefore, it is possible that the overall conformational ensemble is largely dominated by parallel fibrils, with the antiparallel aggregates representing a smaller subset of structures. Recent studies on Aβ mutants from mouse models reveal a similar finding, where a small fraction of fibrillar polymorphs exhibited an antiparallel arrangement(16). Our results underscore the critical significance of obtaining structural information about amyloid aggregates directly from vascular aggregates to complement seeded growth from brain-derived fibrils. Recent studies have revealed similar findings for Aβ plaques, where presence of antiparallel structures in a subset of plaque cores has been demonstrated(32). Interestingly, in recent work, Irizarry et. al. has used IR and Nuclear Magnetic Resonance (NMR) spectroscopy to show that CAA-derived fibrils can have both parallel and antiparallel architecture(29). This agrees with our findings and supports the view that antiparallel fibrils can be concurrently present along with the expected parallel structure in vascular amyloids. Another explanation of the discordance between our results and cryoEM is that the antiparallel structures observed here arise from oligomers and are hence not observed in brain derived fibrils investigated in cryoEM. However, there are no reports that have identified significant accumulation of oligomeric species in blood vessels with CAA progression. Hence, we disregard this possibility and consider the alternative: the antiparallel structure arises from fibrillar aggregates. This leads to the question: what leads to antiparallel fibrils? The other intriguing insight from this work is the potential role of lipids in modulating the structural distribution of aggregates. We find that the relative abundance of lipids in blood vessels correlates specifically to antiparallel β-sheets and not to the overall β-sheet population, which indicates that a. vascular amyloids have a mixed structural ensemble comprised of both parallel and antiparallel aggregates, and b. the formation of the latter may be connected to or mediated by interaction with lipids. The role of lipids in accelerating and modulating the amyloid aggregation pathway is well known(61, 62). Charged lipids surfaces can aid in the nucleation of monomers through electrostatic interactions, thus accelerating the formation of Aβ aggregates and fibrils. Lipid interactions can also influence the structure of Aβ aggregates. While often promoting the formation of the typical parallel β-sheet fibrils, lipids can stabilize distinct oligomeric intermediates, some potentially possessing antiparallel character, which are strongly implicated in membrane disruption and cellular toxicity(63, 64). Amyloid plaques are known to contain lipids colocalized with Aβ aggregates, which further highlights their potential role in altering amyloid aggregation pathways in disease(23, 38). Recent studies using cryo-EM have demonstrated that Aβ40 forms predominantly parallel fibrils in-vitro in presence of 1,2-dimyristoyl-sn-glycero-3-phosphoglycerol (DMPG) vesicles(65). Interestingly, the fibrillar arrangement identified in this work closely resembles that reported by Ghosh and coworkers, who identified an additional outer cross-β layers with antiparallel arrangements in brain-derived fibrils(15). Zhaliazka and coworkers have provided a more nuanced view of potential interactions of different lipids with Aβ and have shown that while most lipids promote higher parallel β-sheet populations, some lipid compositions can lead to an increase in the antiparallel character(66, 67). This suggests that Aβ, when interacting with a heterogeneous mixture of lipids that likely prevail in the brain, could form a mixture of parallel or antiparallel fibrils, or fibrils with mixed β-sheet character.

### Aβ aggregates in parenchyma are structurally different

The common role of Aβ in both CAA and AD has led to the hypothesis that the pathologies of the two diseases can be interlinked. A well-known pathological signature of CAA is the prevalence of dyshoric changes: the spread of Aβ aggregates from blood vessel walls into the surrounding tissue(1–3). Aβ accumulation in blood vessels can hinder its clearance from the tissue, leading to more deposits in the adjoining parenchyma. This can potentially contribute to and affect the pathogenesis of AD. However, it is not known if the same aggregation pathways are prevalent for Aβ in blood vessels and the tissue microenvironment. To understand the structural correlation between vascular and parenchymal Aβ aggregates, we therefore acquired spectra from the Aβ deposits in the tissue adjoining a total of 25 blood vessels, as identified by IHC (Figure 1, also see Figure S3), from two diseased patients corresponding to mild and moderate/severe CAA. The spectra were grouped by blood vessel and projected on to the same basis as those from vascular aggregates to determine the relative contributions of the different secondary structure components. The results of the deconvolution analysis are shown in Figure 5. We observe that the relative population of overall β-sheets increases from mild to moderate/severe CAA, as expected. In contrast, both the antiparallel β-sheet and lipid populations decrease. Furthermore, only the antiparallel population exhibits correlation with lipids (Figure S4). This persistent correlation between the antiparallel population and lipids suggests that the mechanistic pathways adopted by Aβ depends strongly on the chemical environment. As a result, the resulting secondary structure of aggregates can vary significantly between blood vessels and parenchyma, leading to distinctly different disease-specific polymorphs. These findings also provide a rationale for why antiparallel structures have been elusive in cryo-EM studies of brain-derived fibrils: if fibrillar seeds are isolated from the entire parenchyma and not specifically from blood vessels, the dominant structure can be parallel β-sheets, as indicated by our results. In fact, the only study that has reported antiparallel structures in brain-derived fibrils also involved isolating Aβ specifically from vascular aggregates, which provides further credence to this hypothesis and underscores the potential heterogeneity in structure of Aβ fibrils in brain lesions and the need for targeted isolation of seeds for characterization of different pathologies.

**Figure 5.**
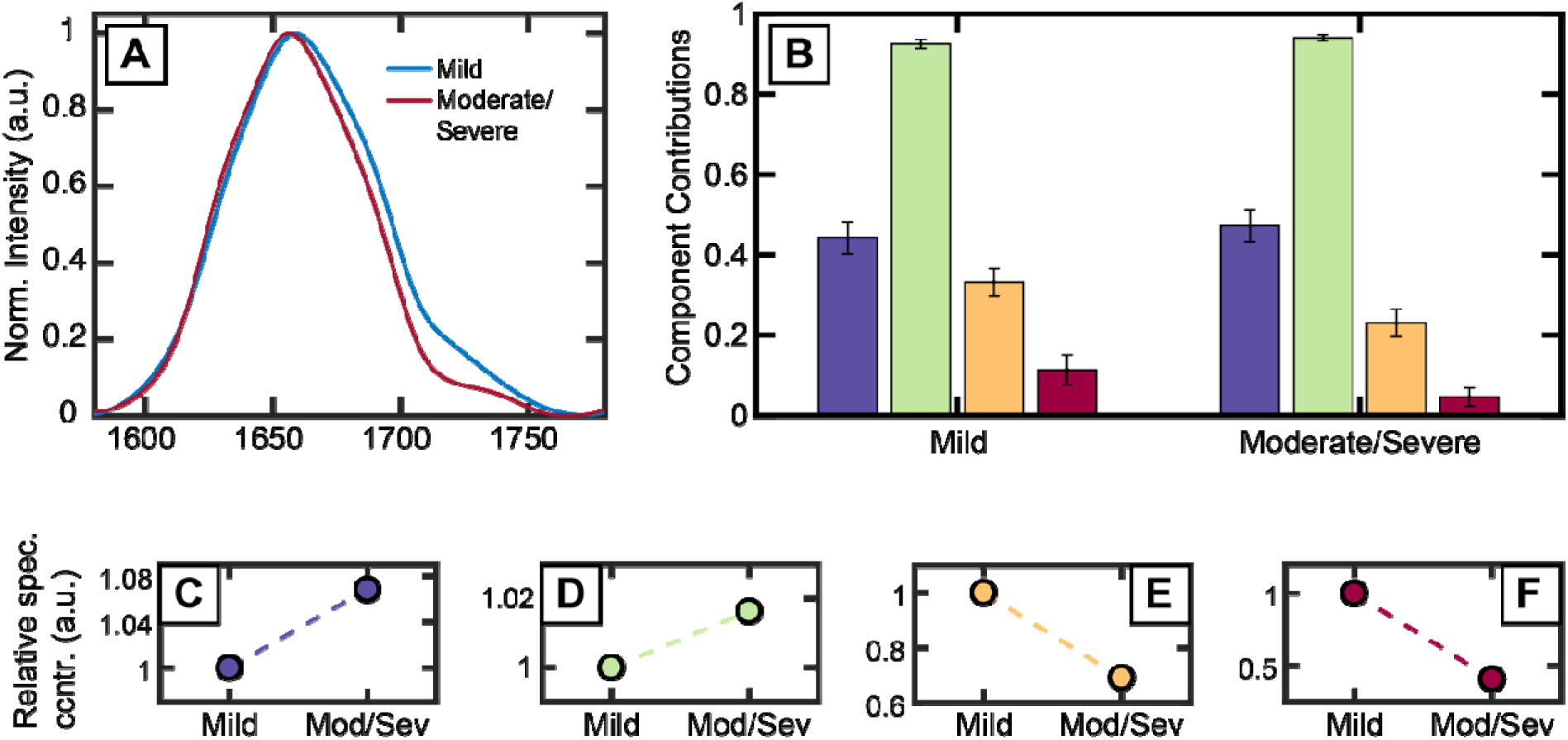
(A) Mean spectra from amyloid deposits in tissue microenvironment adjoining vascular aggregates, as identified by IHC. (B) Contributions of each spectral component determined from deconvolution. (C-F) The relative change in secondary structure populations, as shown in B. The values for each component have been normalized to that of the mild CAA specimens.

### Nanoscale IR spectroscopy validates lipid-induced antiparallel character in Aβ40 aggregates

However, a key caveat needs to be considered here. The results shown above demonstrate the presence of antiparallel β-structure in vascular amyloids; however, these structural fingerprints cannot be uniquely attributed to Aβ or any specific protein from the IR spectra. It is expected from the immunostaining that the spectral features are primarily from Aβ aggregates, but the possibility of other proteins contributing to the antiparallel spectral markers observed here cannot be ruled out. Therefore, to further verify if these antiparallel structures can indeed arise from Aβ and whether they result from lipid interactions, we used nanoscale IR spectroscopy to probe the structure of individual Aβ-40 aggregates. Integrating IR with modalities that offer nanoscale spatial resolution, such as Atomic Force Microscopy, provides a means to characterize the spectra of individual fibrils and the effect of lipids on their structural arrangement. AFM-IR leverages the same photothermal signal as MIP but uses the modulation of the cantilever oscillation for detection, which provides the desired spatial resolution on the scale of individual aggregates(68–70). AFM-IR has been used for characterizing the structure of different amyloid proteins including Aβ under various aggregation conditions(24, 30, 67, 71–73). We focused on answering a specific question relevant to our findings in *ex-vivo* CAA tissues: if/how interaction with a heterogeneous mixture of lipids alters the structure of Aβ fibrils. To that end, we aggregated Aβ-40 in presence of bovine total brain lipid extracts. We used ^13^C Aβ to minimize any spectral overlap between the protein transitions and the lipid. This isotopic substitution is widely used for studying amyloid proteins and redshifts the amide band by ∼30 cm^-1^. Figure 6 A-D show the representative AFM images of Aβ-40 aggregates formed in absence and in presence of lipids after 24 hours of incubation, along with the average AFM-IR spectra acquired from multiple aggregates from different spatial locations. The spectrum from lipids, in absence of any Aβ, is shown in Figure S5. The control Aβ fibrils exhibit a peak at ∼ 1630 cm^-1^, with shoulders at ∼1590 cm^-1^ and ∼ 1660 cm^-1^. This is similar to previously reported AFM-IR spectra of Aβ fibrils(24, 73). Due to the ^13^C amide backbone, the 1590 cm^-1^ band corresponds to overall β-sheets, whereas the 1660 cm^-1^ band represents the antiparallel motif(14, 24, 73). Fibrils formed in presence of lipids exhibit a markedly different spectrum, with a significantly more prominent antiparallel peak, and an additional band at ∼1740 cm^-1^ corresponding to the lipids. The MCR-ALS deconvolution of the spectra, shown in Figure 6E also provides a quantitative validation for this observation, thus unequivocally demonstrating that association with lipids results in elevated antiparallel character in Aβ fibrils. We note that relative distributions of parallel and antiparallel structures in the fibrillar aggregates can evolve with maturation, as has been demonstrated before. Nonetheless, these results provide key evidence in support of the main conclusions drawn from IR spectra of vascular aggregates: that Aβ40 can form antiparallel structures via interactions with lipids, which can have a potential role in neurotoxicity and inflammation in later stages of CAA. Interestingly, the spectra of oligomeric/non-fibrillar species and fibrils (Figure 6) both exhibit elevated antiparallel character. Hence, while these findings provide a molecular basis for observation of antiparallel structures in vascular Aβ aggregates, they do not preclude the possibility of this arising from elevated abundance of oligomeric species in addition to fibrils. Given that oligomeric species are more toxic than mature fibrils, an increase in oligomeric populations in vascular aggregates could have significant implications on the different mechanisms of neurotoxicity associated with CAA progression. However, conclusively attributing the vascular spectra to a specific morphology and/or aggregation stage of Aβ necessitates additional complementary approaches, which are beyond the scope of this report. We hope to address this in future work.

**Figure 6.**
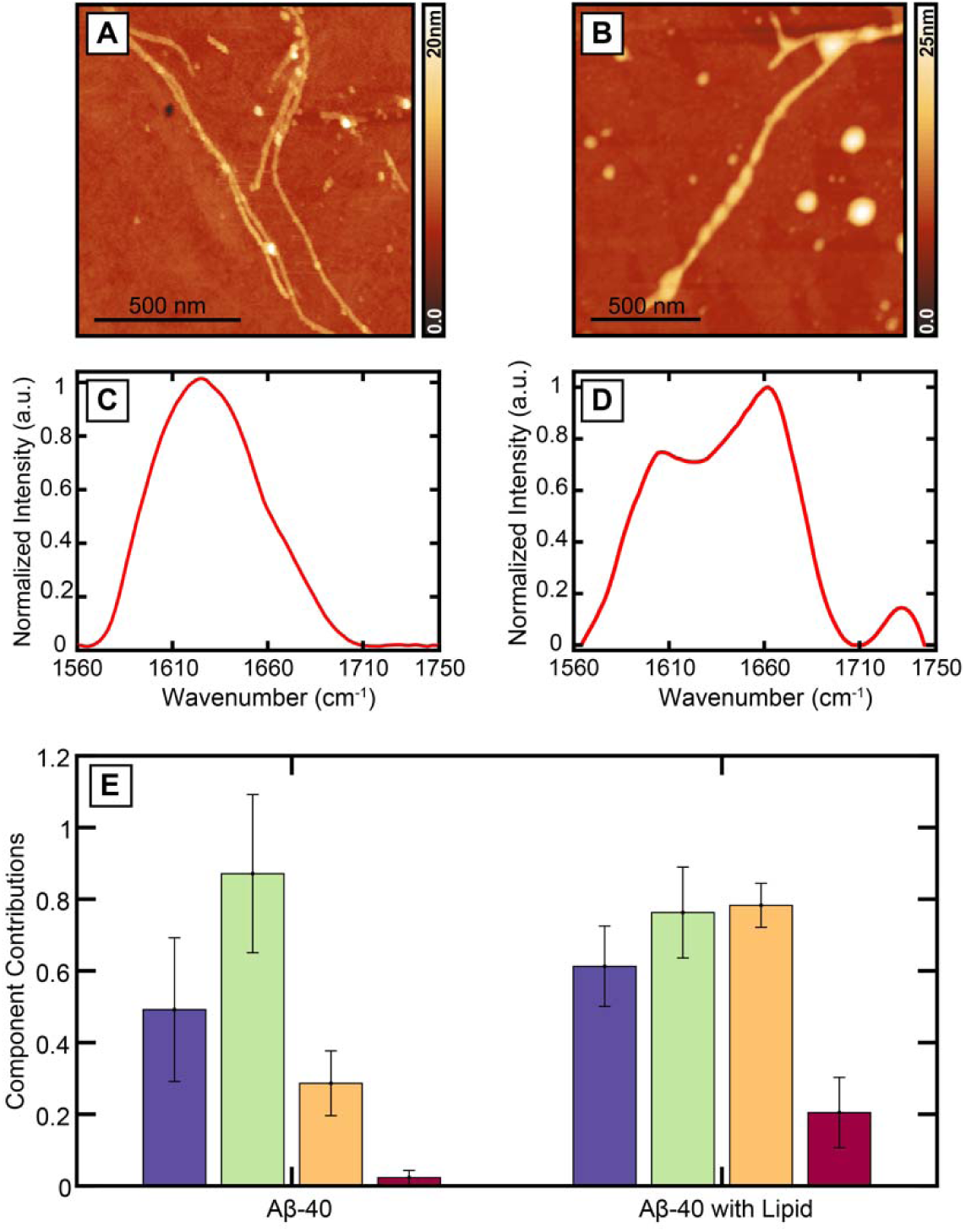
AFM-IR characterization of ^13^C-Aβ40 co-aggregated with brain lipid extract. (A, B) AFM images of ^13^C-Aβ40 and ^13^C-Aβ40 fibrils in presence of brain lipid after 24h of incubation, respectively. (C, D) Average AFM-IR spectra from multiple spatial locations on the aggregates. (E) Spectral deconvolution of AFM-IR spectra into constituent components representing overall β-sheets (blue), random coil (green), antiparallel β-sheets (orange) and lipids (red).

### Conclusions

In summary, the results presented herein paint a unique picture of secondary structure in vascular amyloids which can involve significant populations of antiparallel Αβ fibrils particularly in moderate/severe CAA, resulting from a lipid mediated aggregation pathway. These findings highlight the complexity of amyloid aggregation pathways that may persist in different disease mechanisms and underscore the importance of complementing *in-vitro* studies with direct, *ex-vivo* or *in-vivo* characterizations of amyloid structure. The role of lipids in altering kinetics of amyloid assembly and the structure of aggregates has been demonstrated *in-vitro*; however, observation of disease relevant manifestations of such pathways in the brain are rare. Our work provides a direct validation of the relevance of such interactions in shaping the amyloid landscape in disease. The neurotoxicity of antiparallel aggregates, both oligomeric and fibrillar, is well-established; it is thus possible that their presence in different disease states can open up alternate avenues of neurodegeneration and inflammation. Understanding the different amyloid structures implicated in pathogenesis and the pathways that lead to their formation is critical for the development of targeted therapeutic interventions necessary to address the public health challenge posed by these protein misfolding diseases.

## Acknowledgement

This work was supported by the National Institutes of Health (Award R35 GM138162 to A. G.). The Duke-UNC ADRC is supported by the National Institutes on Aging (P30 AG072958). The content is solely the responsibility of the authors and does not necessarily represent the official views of the National Institutes of Health.

## Notes

### Competing Interest Statement

The authors have declared no competing interest.

