## Supporting Information for "Lipid Mediated Formation of Antiparallel Aggregates in Cerebral Amyloid Angiopathy"

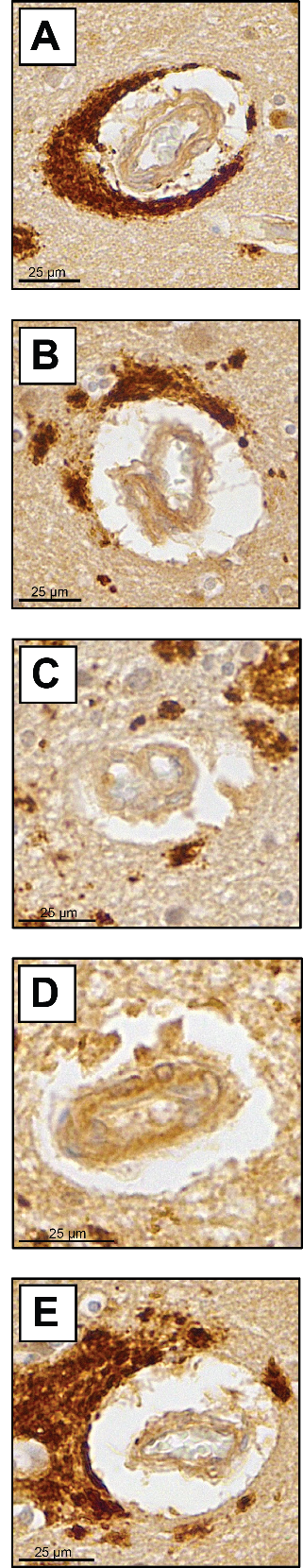

**Figure S1.** (A-E) Brightfield optical images of vascular amyloid deposits from mild CAA case. Scale bar is 25 μm.

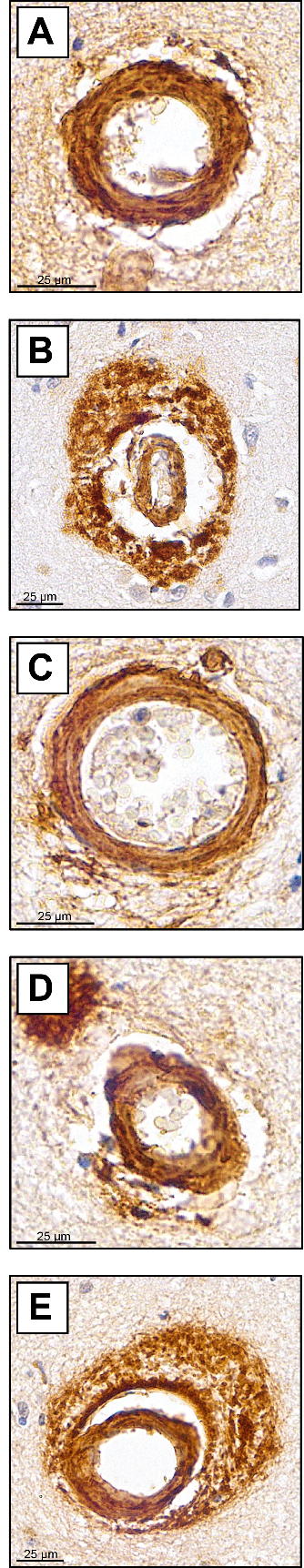

**Figure S2.** (A-E) Brightfield optical images of vascular amyloid deposits from severe CAA case. Scale bar is 25 μm.

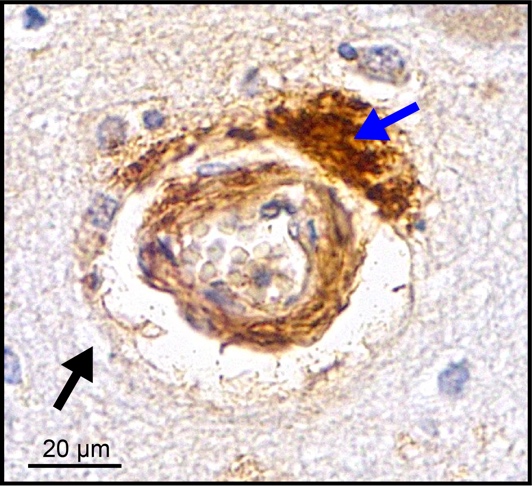

**Figure S3.** Representative brightfield image of vascular aggregate indicating presence of parenchymal amyloid aggregates in the tissue microenvironment (blue arrow). The black arrow indicates a representative spatial location from which control, amyloid-negative spectra are acquired.

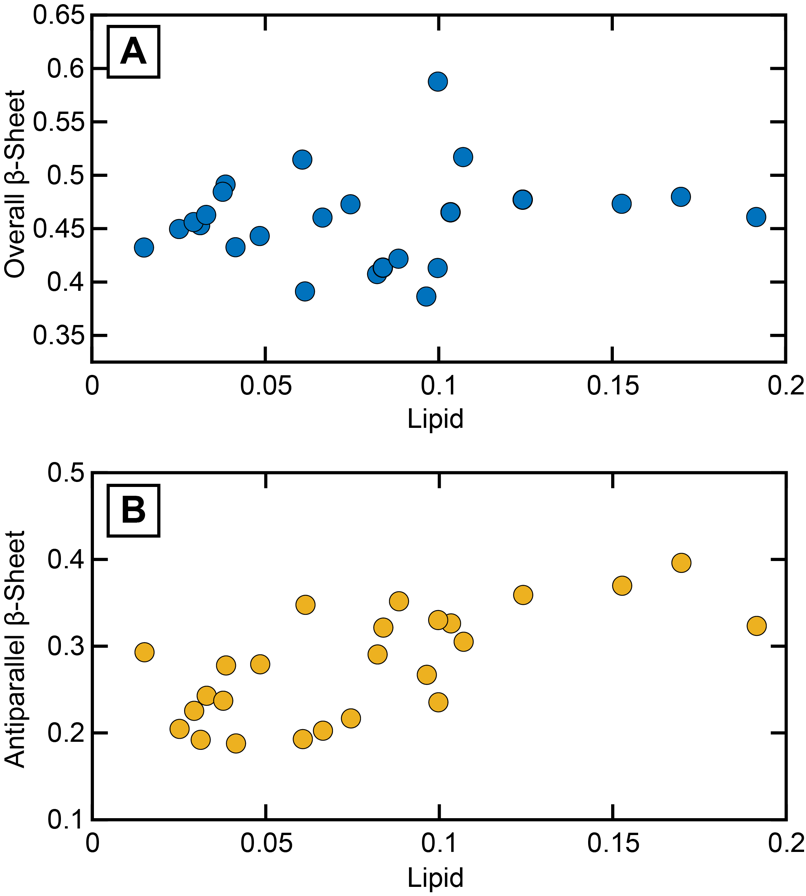

**Figure S4.** Scatter plot of populations of (A) overall β-sheets and (B) antiparallel β-sheets vs lipids in parenchymal amyloid deposits, as determined from spectral deconvolution using MCR-ALS.

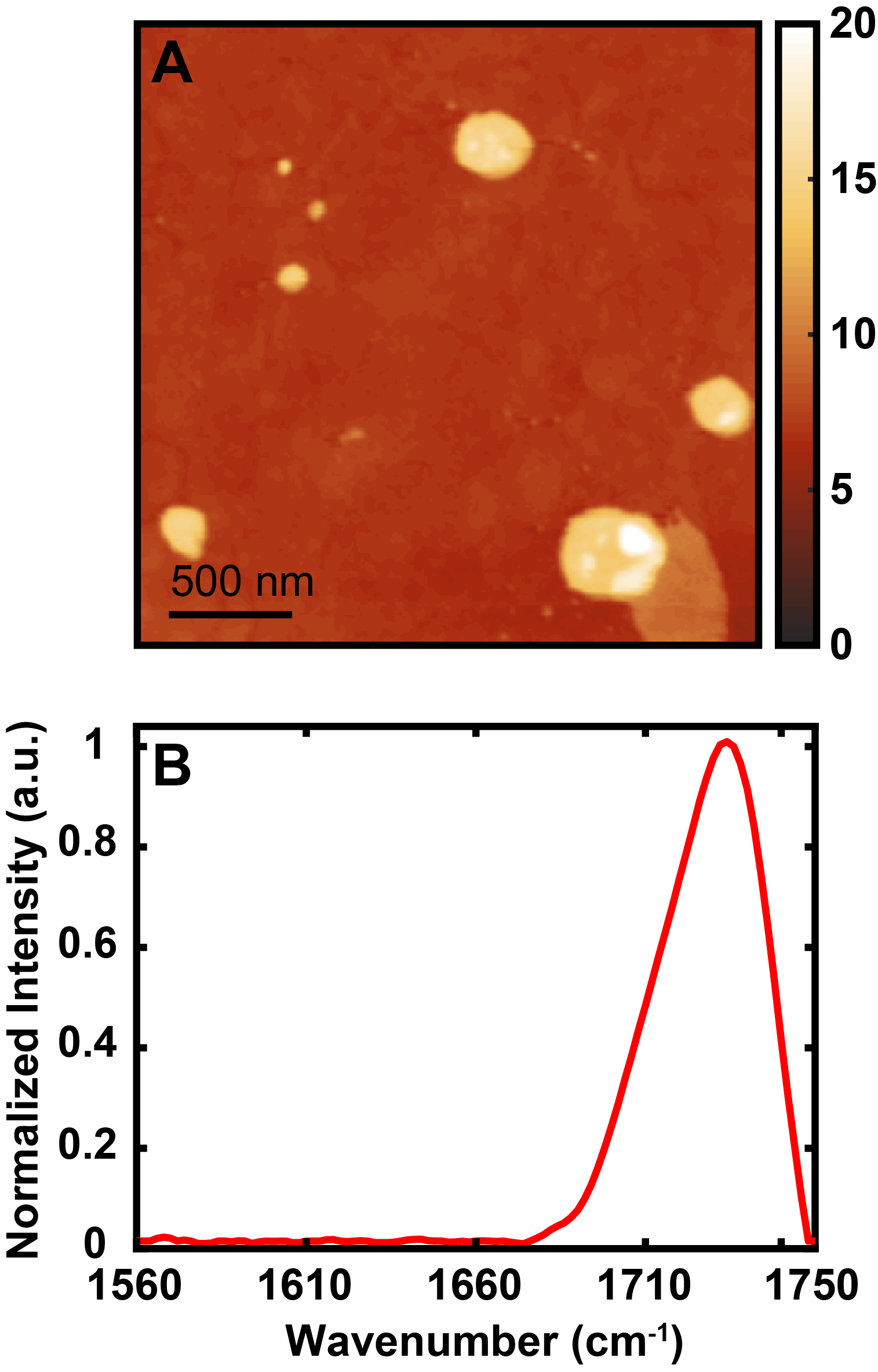

**Figure S5.** AFM-IR characterization of vesicles of brain lipid extract in 10 mM phosphate buffer. (A) AFM topographic image after 24h of incubation. (B) Average IR spectra different spatial locations in the AFM image.

| **Age** | **Sex** | **Neuropathological Diagnoses** | **Number of blood vessels studied** |
| --- | --- | --- | --- |
| 85  (Control) | F | ADNC- Not  A = 0 (Thal 0) \| B = 1 (Braak Stage II) \| C = 0 \| | 20 |
| 76  (mild CAA) | F | ADNC – Intermediate  A = 3 (Thal 4) \| B = 2 (Braak Stage III) \| C = 2 | 20 |
| 80  (moderate/severe CAA) | F | ADNC - Intermediate  A = 3 (Thal 5) \| B = 2 (Braak Stage IV) \| C = 2 | 20 |
| 81  (moderate/severe CAA) | F | ADNC - High  A = 3 (Thal 5) \| B = 3 (Braak Stage V) \| C = 2 | 10 |
| 77  (mild CAA) | F | AD NPath Changes – Intermediate  A = 2 (Thal 3) \| B = 2 (Braak Stage III) \| C = 0 | 10 |

**Table S1.** Tissue specimens used in this study and corresponding neuropathological diagnoses. The CAA designations are based on IHC staining.

| **Parallel β-sheets** | **p-value** |
| --- | --- |
| Control vs Mild CAA | <0.0001 |
| Control vs Moderate/Severe CAA | <0.00001 |
| Mild vs Moderate/Severe CAA | <0.00001 |

| **Antiparallel β-sheets** | **p-value** |
| --- | --- |
| Control vs Mild CAA | 0.33 |
| Control vs Moderate/Severe CAA | 0.007 |
| Mild vs Moderate/Severe CAA | 0.12 |

| **Lipids** | **p-value** |
| --- | --- |
| Control vs Mild CAA | <0.00001 |
| Control vs Moderate/Severe CAA | <0.00001 |
| Mild vs Moderate/Severe CAA | 0.09 |

**Table S2 .** Results from statistical analysis of secondary structure contributions to spectra, as determined by MCR-ALS. The statistical significance was determined by performing Welch’s ANOVA test, followed by Games-Howell post-hoc test, to account for inequal data sizes and variances between the groups.
